# Mouse model of multiple sclerosis induced by disrupting vesicular transport in oligodendrocytes

**DOI:** 10.1101/2023.08.24.554669

**Authors:** Chun Hin Chow, Mengjia Huang, Jayant Rai, Hidekiyo Harada, Sarah Eide, Hong-Shuo Sun, Zhong-Ping Feng, Philippe P. Monnier, Kenichi Okamoto, Liang Zhang, Shuzo Sugita

## Abstract

Multiple Sclerosis is an autoimmune demyelination disorder with unknown etiology. Despite the myelin damage, the roles of myelinating oligodendrocytes in driving disease progression remain unknown. We hypothesize that disrupting vesicular transport in oligodendrocytes during adolescence will disrupt myelin integrity and causes neuroinflammation. By creating a mouse model of SNAP-23 conditional knockout in mature oligodendrocytes, we showed that impairment in vesicular trafficking in oligodendrocytes causes demyelination. Neuroinflammation with infiltration of peripheral immune T cells into the central nervous system was observed accompanied by demyelination. Mechanistically, SNAP-23 removal in oligodendrocytes caused abnormal axon-myelin structures and impaired myelin protein trafficking, both can contribute to autoimmune activation and demyelination. With our novel animal model, we propose that oligodendrocyte injury is an endogenous early event in triggering Multiple Sclerosis.

**One-Sentence Summary:** Impaired vesicular transport in oligodendrocytes in adults caused demyelination and inflammation driving Multiple Sclerosis

## Main Text

Cures for Multiple Sclerosis (MS), a demyelination disorder caused by an autoimmune attack on myelin, are lacking due to an insufficient understanding of the etiology (*1, 2*). Animal research on MS often utilizes the experimental autoimmune encephalomyelitis (EAE), an “outside-in” model, that triggers demyelination by externally driven autoimmunity (*2-5*). To account for the endogenous pathophysiology, the “inside-out” model which considers myelin and the myelinating glia oligodendrocytes as the primary injury is proposed (*2, 4*). The presence of disease-associated oligodendrocytes in MS models further indicates oligodendrocytes as a crucial component in MS pathology (*6-8*). However, the mechanism of oligodendrocytes leading to demyelination and autoimmunity remains unknown.

Myelin is not a stable lipidic ensheathment. Lipids and proteins are continuously synthesized and delivered from oligodendrocytes soma to myelin (*9-13*). Vesicular transport is the basis of myelin materials transport (*10, 14, 15*). Soluble N-ethylmaleimide-Sensitive Factor Attachment Proteins Receptor (SNARE) proteins which mediate vesicular fusion are responsible for trafficking myelin proteins (*15-19*). During development, vesicular SNAREs VAMP2/3 are necessary for myelin formation by modulating protein insertion to myelin and oligodendrocyte precursor cells (OPC) maturation (*18, 20*). However, MS is an adult-onset disorder, and the role of SNARE-mediated fusion in mature oligodendrocytes remains elusive. Moreover, the target-SNARE SNAP isoform is not identified. Oligodendrocytes express SNAP-23 instead of the conventional neuronal SNAP-25 isoform (*14*). Therefore, we hypothesize that SNAP-23 mediated vesicular fusion transports myelin proteins to myelin, and disruption of such process will impair myelin integrity and trigger autoimmune attacks on myelin in adults.

Here, we showed that the removal of SNAP-23 in mature myelinating oligodendrocytes of adolescent mice triggered demyelination in adults. Neuroinflammation was accompanied by peripheral T cell infiltration into the central nervous system (CNS). Lastly, we demonstrated that impaired vesicular fusion machinery altered axon-myelin structure and myelin glycoproteins trafficking, which (potentially) serve as the basis for myelin damage and T cell activation.

### SNAP-23 removal in mature oligodendrocytes causes demyelination in adults

To investigate the impact of damaging vesicular transport in oligodendrocytes, we generated the *Plp1-CreERT; SNAP-23flox/flox* mice (SNAP-23 icKO). Tamoxifen injection at 3-to-4-week-old mice knocked out SNAP-23 in oligodendrocytes (Fig. 1A). Genotyping by polymerase chain reaction (PCR) of the spinal cord after tamoxifen injection showed the recombinant band in the SNAP-23 icKO but not in the *SNAP-23flox/flox* (Ctrl), indicative of successful knockout (Fig. 1B, C). To evaluate the demyelination phenotype, we employed a modified EAE scoring system to evaluate the mice weekly post tamoxifen injection (pti) (*12*). We observed the development of tremors, limp tail, hindlimb weakness, and paresis with a peak of clinical scores at 5-to-10-week pti (Fig. 1D, E). A significant decline in the weight of males and females was observed in the peak demyelination phase (Fig. S1). At 4 to 5 weeks pti, SNAP-23 icKO was falling significantly earlier compared to the control throughout the four-day accelerating rotarod test (Fig. 1F). The poor rotarod performance reflected demyelination-induced motor deficits.

**Fig. 1.**
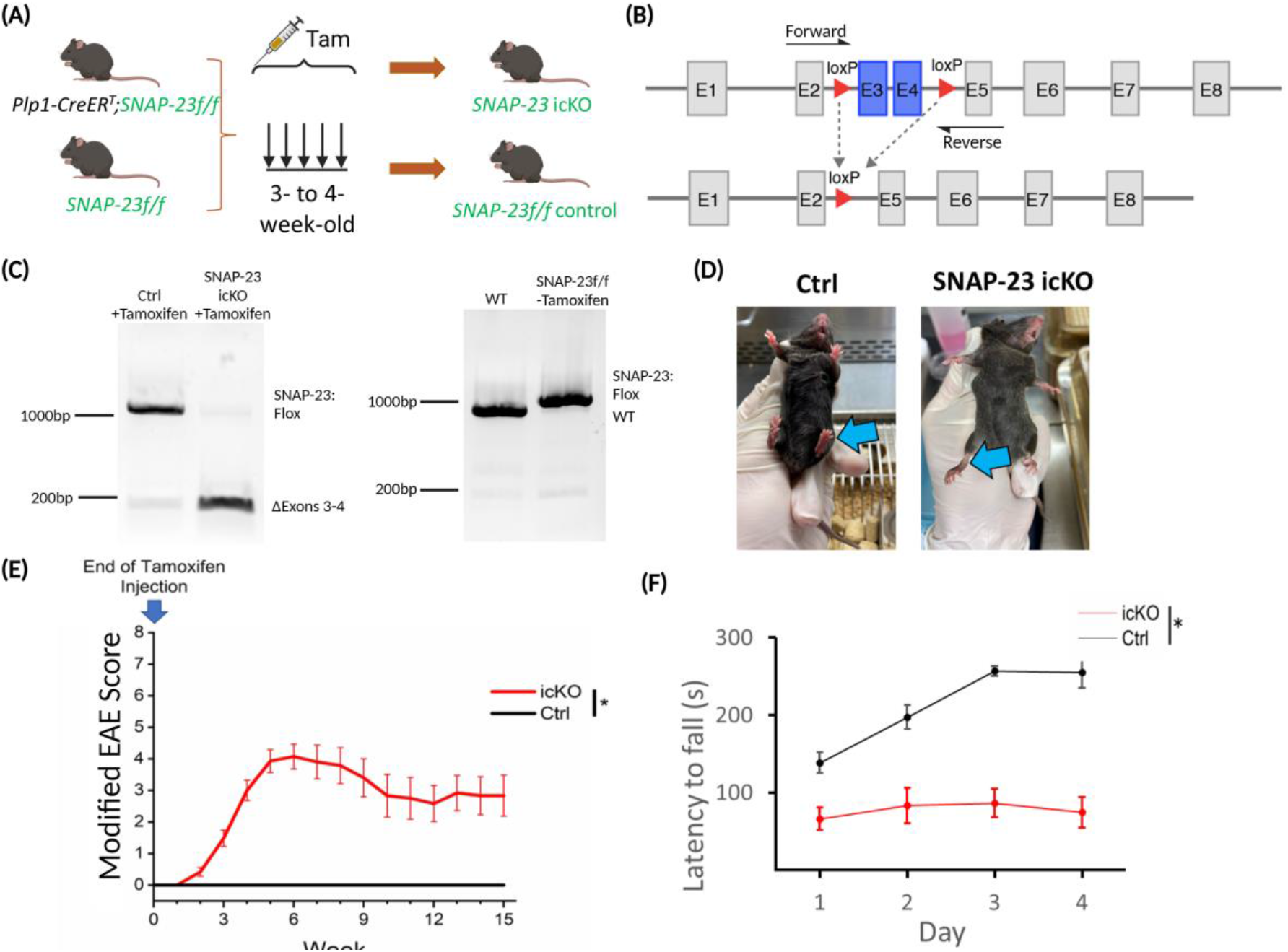
Removal of SNAP-23 in mature oligodendrocytes caused demyelination in adults. **(A)** Schematic of the knockout strategy and timeline. Tamoxifen (100mg/kg) was applied to 3- to-4-week-old mice to induce knockout (SNAP-23 icKO and SNAP-23f/f (Ctrl)). **(B)** *SNAP-23* gene structure. Exons 3 and 4 (E3 and E4) are removed in the knockout. Forward and Reverse primers for genotyping are indicated. **(C)** PCR spinal cord from ctrl and SNAP-23 icKO after tamoxifen injection confirming knockout of SNAP-23 (left). PCR from tail samples of wildtype (WT) and *SNAP-23f/f* control animals. **(D)** Example of mice showing hindlimb weakness in the SNAP-23 icKO which hindlimbs point down instead of upward (blue arrow). **(E)** Weekly progression of modified EAE scores qualifying tremor, limp tail, hindlimb weakness, and paresis. Each score has a scale of 0-2, 2 being the most severe. Control (Ctrl): n=10-22; iCKO: n=12-31. *p<0.05 by 2-way ANOVA followed by Tukey post hoc test. **(F)** Accelerating rotarod on animals 4 to 5 weeks pti across 4 days. Ctrl: n=8; iCKO: n=10. *p<0.05 by Mixed ANOVA. Data are mean±SEM.

To verify the demyelination phenotype, we performed structural and ultrastructural analyses on CNS white matter on 5 to 7 weeks pti. Luxol fast blue (LFB) staining indicated a reduction of myelin in the optic nerves, spinal cord, and brain (Fig. 2A, B, S2A). Independently, we verified the loss of myelin with Fluoromyelin staining in the CNS (Fig. S2B). Concurrently, we imaged the optic nerve cross sections by transmission electron microscopy (TEM). SNAP-23 icKO optic nerves had a significant reduction in the density of myelinated axons compared to the control (Fig. 2C, D). Smaller myelinated axons with sizes (0.3-0.6μm) were lost in the SNAP-23 icKO (Fig. 2E). To evaluate the myelin sheath thickness, we quantified the G-ratio, which is the ratio of total myelinated axon diameter over axon diameter. SNAP-23 icKO optic nerves contained axons with a larger G-ratio, indicating a thinner myelin ensheathment compared to the control (Fig. 2F). These structural abnormalities together pointed towards demyelination that happens in the adult SNAP-23 icKO mice across the CNS.

**Fig. 2.**
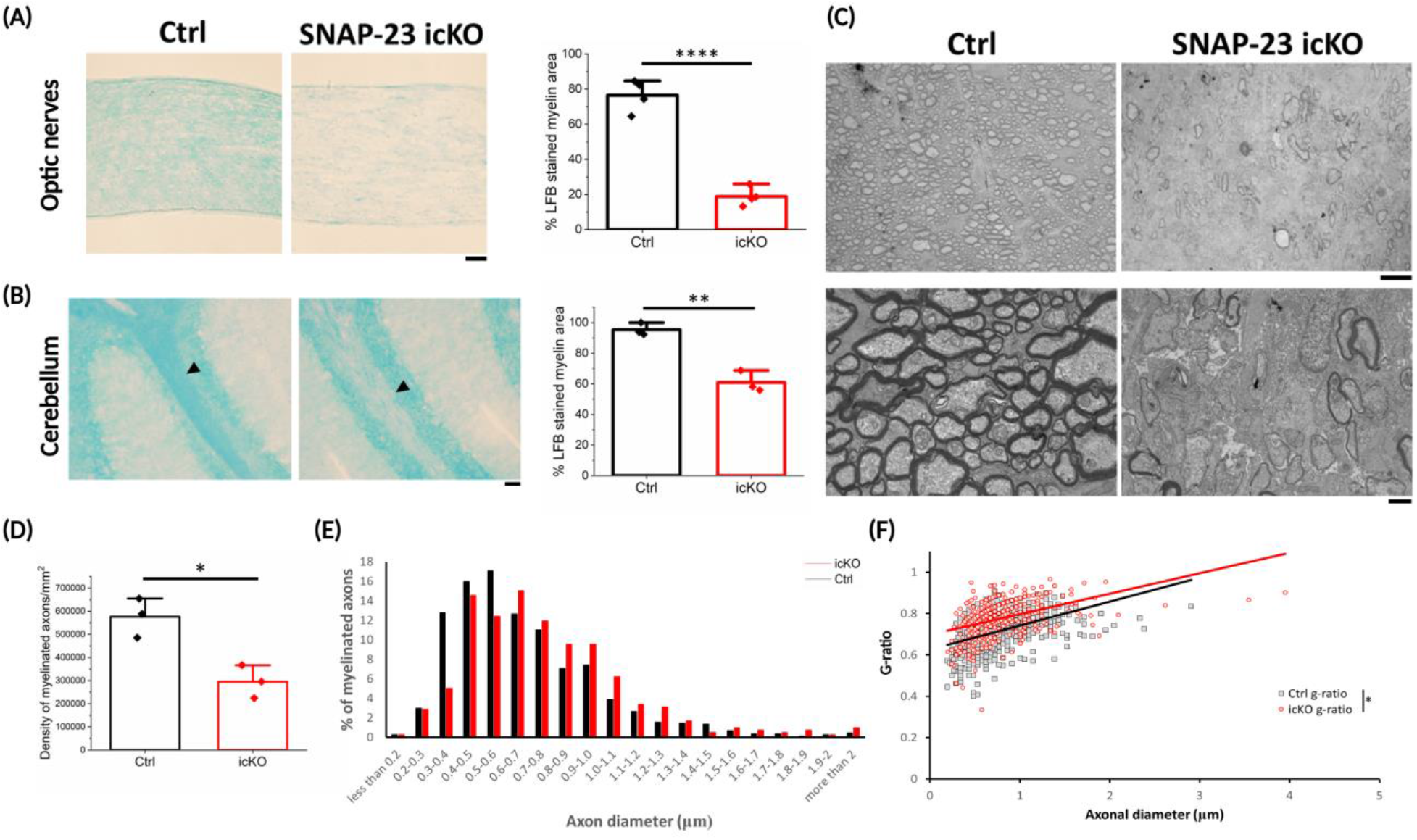
Structural demyelination in the CNS of SNAP-23 icKO. **(A)** LFB staining of the optic nerves and quantification of LFB+ myelin area in the optic nerves. n=4 optic nerve sections from 2 animals in each group. **(B)** LFB staining of the cerebellum. Quantification of LFB+ myelin area in the cerebellum. n=3 brain sections in each group. Scale bar: 50μm. **p<0.01, ****p<0.0001 by two-sample t-test. Data are mean±SEM. **(C)** Ultrastructure of optic nerves cross-section. Scale bar: 10μm (top), 1μm (bottom). **(D)** The density of myelinated axons was decreased in the SNAP-23 icKO. *p<0.05 by two-sample t-test. **(E)** Relative distribution of myelinated axons of different sizes. **(F)** G-ratio quantification of the myelin sheath. *p<0.05 by simple linear regression analysis of intercepts. n=3 optic nerve sections from 2 animals in each group.

### Functional deficits in nerve conduction in SNAP-23 icKO

To investigate the functional consequences of demyelination in our model, we measured the conductance of the optic nerves by *ex vivo* recording of compound action potential (CAP). In the control, optic nerves showed a CAP profile with three peaks, P1, P2, and P3 corresponding to the largest to smallest myelinated axons. However, CAP in the SNAP-23 icKO 5 weeks pti had a delayed P1 and P2 with an altered P3 waveform (Fig. 3A). Latencies to peaks in the SNAP-23 icKO were significantly larger compared to the control (Fig. 3B). The amplitude of each peak was smaller in the SNAP-23 icKO while the stimulus artefact remained similar between the SNAP-23 icKO and control (Fig. 3C, E). The full width at half-maximum (FWHM) of CAP of SNAP-23 icKO was significantly larger than that of the control, indicating wider waveforms with altered synchrony in nerve conduction (Fig. 3D).

**Fig. 3.**
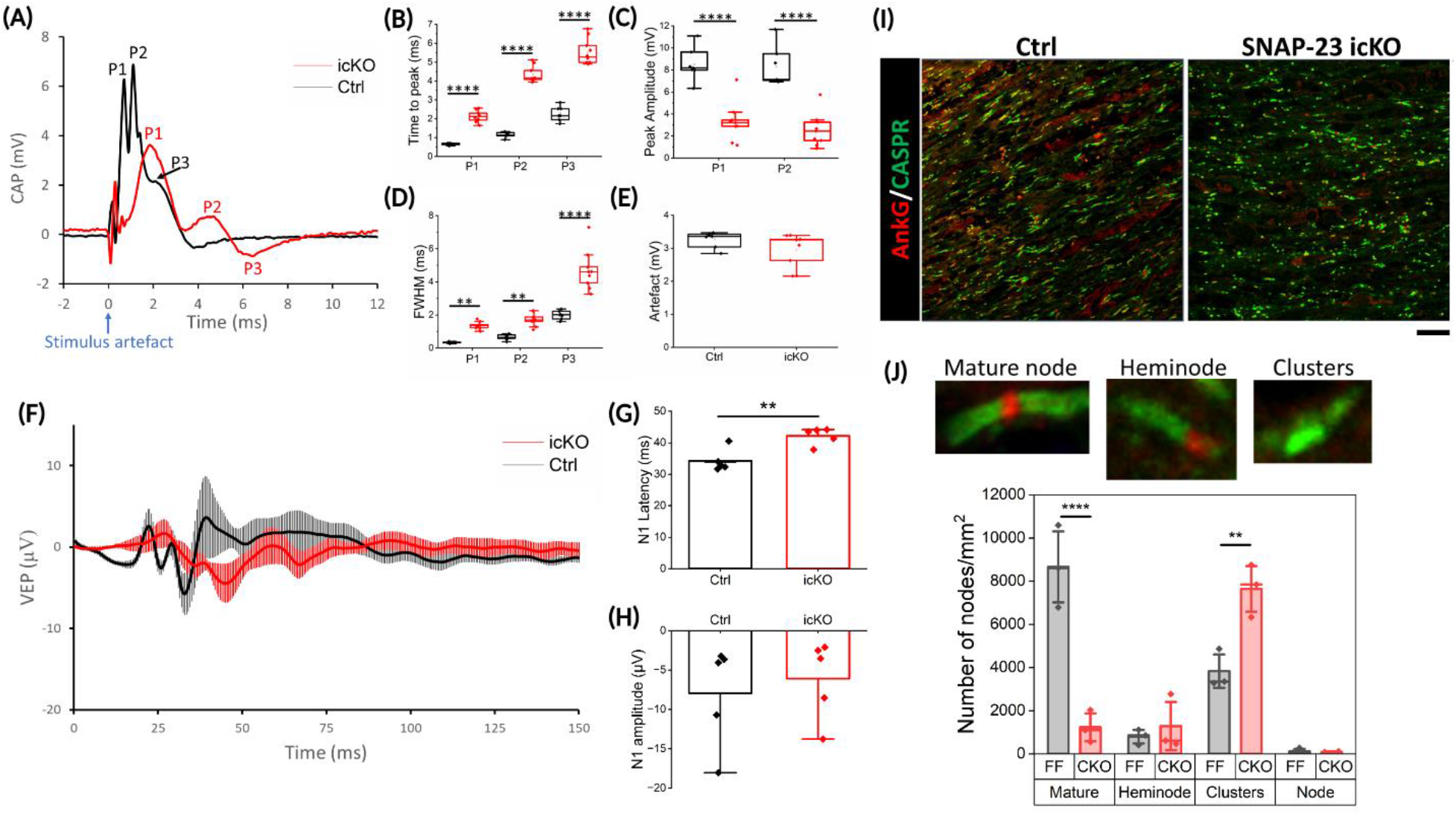
Impaired functional conduction with altered nodes of Ranvier in SNAP-23 icKO. **(A)** Example traces of *ex vivo* compound action potential (CAP) measurement in the optic nerves. **(B)** Latency to peaks. **(C)** The peak amplitude of P1 and P2. **(D)** The full width at half-maximum (FWHM) of each peak. **(E)** Stimulus artefact amplitude. Ctrl: n=7 nerves from 5 animals; iCKO: n=10 nerves from 5 animals. **p<0.01, ****p<0.0001 by One-way ANOVA with Tukey post hoc test. **(F)** Average traces of Ctrl (black) and SNAP-23 icKO (red) visual evoked potential (VEP) at peak demyelination stage (5-to-6-week pti). **(G)** Latency of N1. **(H)** Amplitude of N1. n=5 in each group. Data are mean±SEM. *: p<0.05 by two-sample t-test. **(I)** Ankyrin G (AnkG, nodal protein in red), CASPR (paranode protein in green). Scale bar: 20μm. **(J)** Quantification of the number of mature nodes, heminodes, paranode clusters, and nodes only. Examples of a mature node, heminode, and cluster are shown at the bottom. n=3 from each group. *p<0.05, ****p<0.0001 by One-way ANOVA with Tukey post hoc test.

Beyond *ex vivo* electrophysiology measurement, we recorded *in vivo* visual evoked potential (VEP) as a measurement of signal conduction along the visual pathway (*21*). SNAP-23 icKO 5 to 6 weeks pti showed a significant delay in N1 latency compared to the control (Fig. 3F, G, S3). Yet, the amplitude was not significantly different (Fig. 3H). Hence, impaired SNAP-23-dependent vesicular trafficking in oligodendrocytes impaired visual signal conduction.

Action potential generation and propagation in myelinated axons happen at the nodes of Ranvier. We hypothesized that demyelination induced by SNAP-23 icKO will alter the arrangement of the nodes and lead to conduction deficits. To visualize the nodes of Ranvier in the optic nerves, we performed immunostaining of Ankyrin G (AnkG) and contactin associated protein 1 (CAPSR), which are located at the node and paranodes respectively (Fig. 3I). A mature node of Ranvier consists of a node flanked by two paranodes, while improper nodal arrangements include heminode (AnkG with CASPR on one side only), paranodes clusters, and node only (*22, 23*). While the control optic nerves comprised mainly mature nodes, the SNAP-23 icKO had a decrease in the number of mature nodes and an increase in CASPR-only paranode clusters (Fig. 3J). Hence, demyelination induced by impaired oligodendrocytes trafficking disrupted the nodes of Ranvier, leading to decreased sensory signal conduction speed and impaired synchrony in axonal conduction.

### SNAP-23 icKO caused neuroinflammation and T cells infiltration into the CNS

An important question in MS is the emergence of inflammation and autoimmune response attacking the myelin. Most studies inducing apoptosis of oligodendrocytes failed to induce T cell activation (*9, 24-26*). Thus, we hypothesized that rather than cell death, the disrupted oligodendrocytes functions and myelin damage trigger inflammatory responses. We demonstrated that by 5 weeks pti, there was an upregulation in astrocytes, by GFAP staining in the corpus callosum, cerebellum, spinal cord, and optic nerves (Fig. 4A, B, S4A-C). Microglia density, indicated by IBA-1 staining was also increased in the CNS (Fig. 4C, D, S4D-F). Thus, SNAP-23 icKO induced neuroinflammation with CNS resident glial cells. Further, we detected the presence of CD3+ T cells in the cerebellum and the spinal cord white matter of the SNAP-23 icKO but not in the control mice (Fig. 4E, S5). Therefore, oligodendrocyte dysfunctions triggered activation and infiltration of T cells into the CNS parenchyma.

**Fig. 4.**
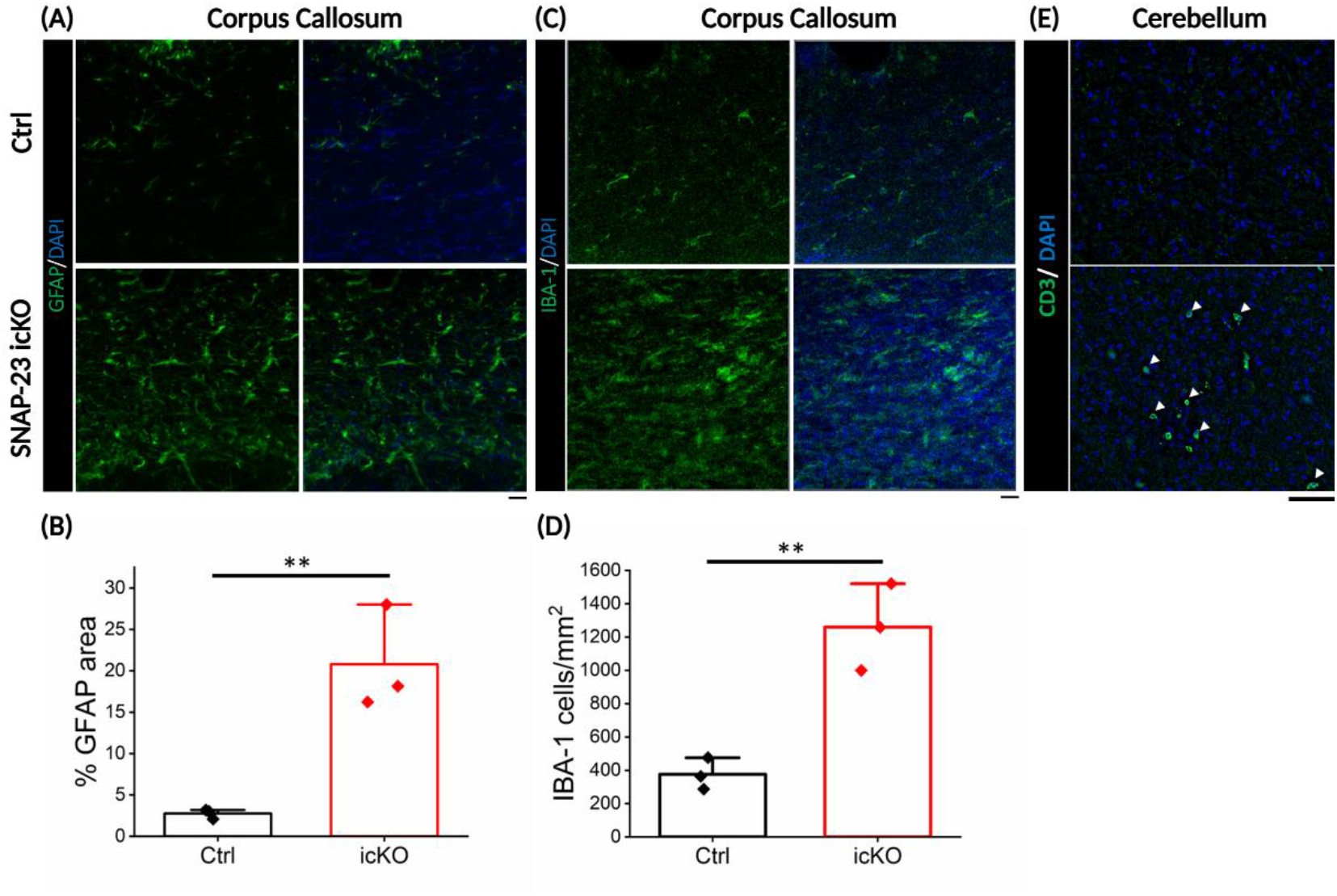
Neuroinflammation and T cell infiltration into the CNS in the SNAP-23 icKO. **(A)** GFAP staining of astrocytes in the corpus callosum. **(B)** Quantification of GFAP+ area in (A). **(C, D)** IBA-1 staining and quantification of microglia density in the corpus callosum. n=3 in each group. Data are mean±SEM. **: p<0.01 by two-sample t-test. **(E)** CD3 staining in the cerebellum. CD3+ cells are evident in the SNAP-23 icKO. Scale bar: 20μm.

### Vesicles accumulation in the myelin and abnormal axonal structure caused by SNAP-23 icKO

Vesicular fusion events happen at the inner tongue of the myelin sheath for adding new myelin materials (*10*). We hypothesize that the removal of SNAP-23 will block vesicular fusion and leads to the accumulation of vesicles at the inner tongue affecting axon-myelin integrity. Through the optic nerves ultrastructural analysis, we found myelinated axons with enlarged inner tongues and an accumulation of vesicles inside (Fig. 5A, B). The insignificant increase can be accounted for by the loss of most myelinated axons at this stage (Fig. 2D). Furthermore, we observed other myelin and axonal abnormalities in the SNAP-23 icKO. Axonal swelling and spheroids, where axons are accumulated with endolysosomal vesicles indicating impaired axonal transport (*27-29*), were present in the SNAP-23 icKO (Fig. 5C). Myelin outfoldings, which suggest the presence of excess myelin and inefficient myelination (*30*), were evident in the SNAP-23 icKO (Fig. 5D). Hence, impairment of vesicular fusion to the myelin sheath and inadequate support to axons were two underlying factors contributing to the demyelination and functional deficits observed in the SNAP-23 icKO mice.

**Fig. 5.**
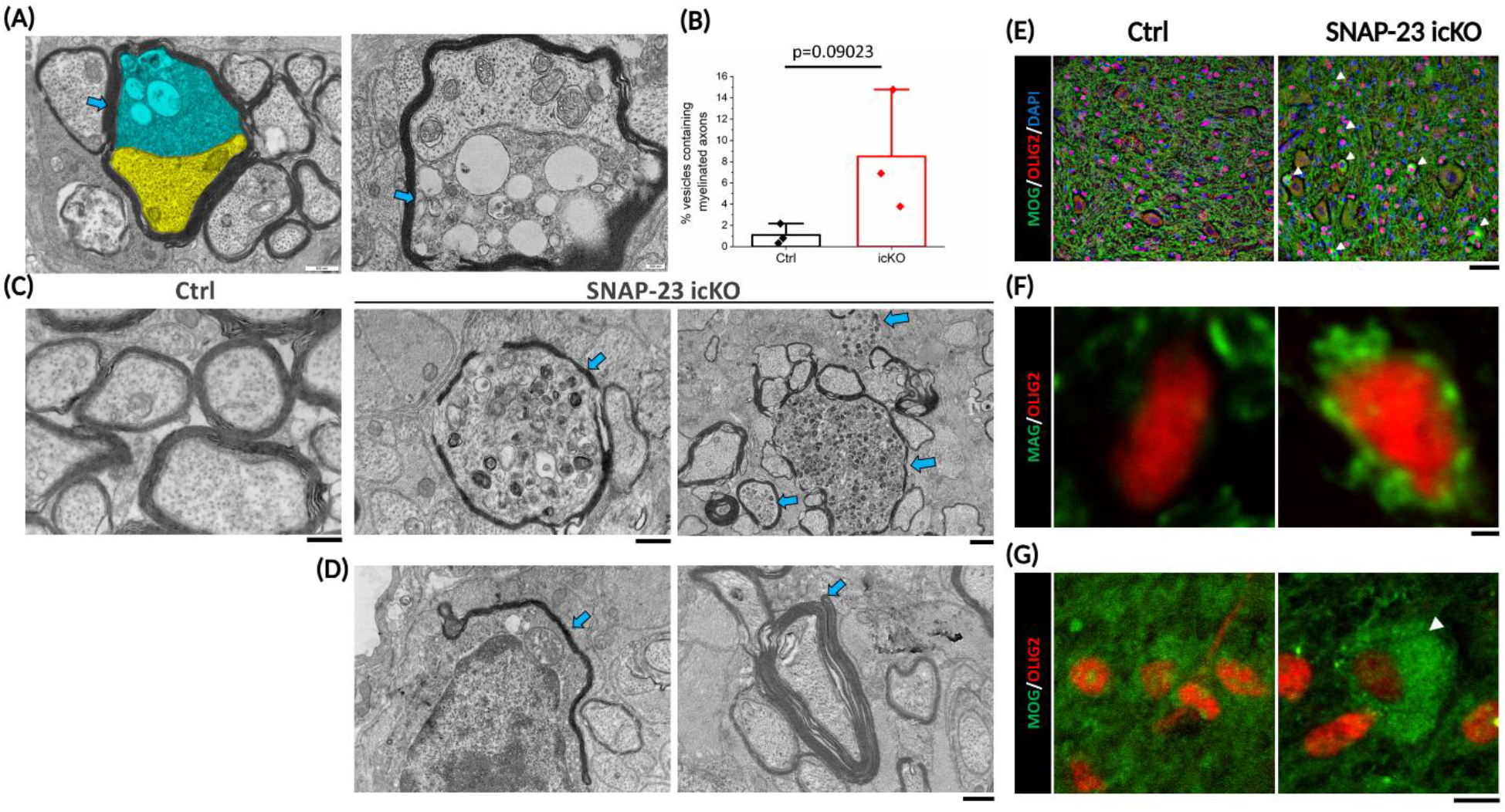
Impaired vesicular fusion and protein trafficking induced by SNAP-23 removal in oligodendrocytes. **(A)** Examples showing vesicle accumulation (arrows) at the inner tongue (blue) adjacent to the axon (yellow). **(B)** Percentage of myelinated axons with vesicle accumulation. n=3 optic nerve sections from 2 animals in each group. p=0.09023 by two-sample t-test. **(C)** Presence of axonal spheroids in SNAP-23 icKO. TEM of the optic nerves showed normal myelinated axons in the control. Axonal swelling and spheroid structures were observed in SNAP-23 icKO (arrows). Scale bar: 400nm in the left two images, 1μm in the right image. **(D)** Myelin outfoldings were present in the SNAP-23 icKO optic nerves (arrows). Scale bar: 500nm. **(E)** MOG staining with OLIG2, oligodendrocyte lineage cell marker, and DAPI in the spinal cord. Arrowheads point to oligodendrocytes with perinuclear localization of MOG. Scale bar: 40μm. **(F)** Two-photon imaging of MAG (Scale bar: 2μm) and **(G)** MOG (Scale bar: 10 μm) with OLIG2 in the spinal cord. Perinuclear localization or proteins was observed (arrowhead).

### Protein trafficking was disrupted in SNAP-23 icKO oligodendrocytes

One of the key functions of SNARE proteins is protein trafficking, in which transmembrane proteins are inserted into the cell membrane via vesicular fusion. During development, VAMP-2/3-mediated exocytosis incorporates axon-myelin adhesion proteins into the myelin sheath (*18*). Given the ability of VAMP-3 to form a SNARE complex with SNAP-23 (*14*), we hypothesized that disruption of SNAP-23 will impair myelin protein trafficking. We performed immunohistochemistry on myelin basic protein (MBP), proteolipid protein (PLP), myelin-associated glycoprotein (MAG), and myelin oligodendrocyte glycoprotein (MOG). In the cerebellum, there were no changes in the staining pattern of MBP and PLP, but we observe punctate rather than uniform staining for MAG (Fig. S6). For MOG, there was a striking perinuclear staining observed in the spinal cord (Fig. 5E). To further investigate protein mislocalization, we utilized two-photon microscopy to visualize protein distribution around the oligodendrocyte nucleus (Olig2+) in the spinal cord. Both MAG and MOG demonstrated a nuclear surrounding staining pattern that was not present in the control (Fig. 5F, G). MOG revealed a clear perinuclear cytoplasmic staining suggesting the improper localization of MOG (Fig 5G). Therefore, the inability of oligodendrocytes to transport critical myelin proteins was one of the mechanisms driving demyelination and inflammation in the CNS.

### Partial remyelination in SNAP-23 icKO

Relapsing-remitting MS (RRMS) involves a recovery (remit) period of symptoms where endogenous remyelination potentially underlies the repairment (*31*). Firstly, SNAP-23 icKO mice had their weight recovered past the peak demyelination (Fig. S1). Modified EAE scores significantly decreased at 6 months compared to 6 weeks dpi (Fig. S7A). Our VEP data on SNAP-23 icKO mice 2 to 7 months pti showed no difference in N1 latency and amplitude with the control (Fig. S7B-D). Therefore, endogenous remyelination, although incomplete, occurred after initial oligodendrocytes damage. Unexpectedly, tracking the mice longitudinally, 11 out of 16 of SNAP-23 icKO fur color turned white (Fig. S7E), which hinted at potential myelin damage in the peripheral nervous system (*32*).

In summary, adult myelin requires continuous replenishment of lipids and proteins to maintain its structure (*9-11, 13*). Here, we demonstrated that removing SNAP-23 in oligodendrocytes causes demyelination with neuroinflammation and axonal pathologies. Our model, therefore, pointed to a previously underrated role of oligodendrocytes in MS progression.

To date, the etiology and development of autoimmunity remain unclear. “Outside-in” animal models including EAE have been deemed unnatural in MS development and provide minimal insights into disease progression (*2, 4*). A recent study highlighted Epstein-Barr virus (EBV) as the major risk leading to MS (*33*). Although EBV altered the immune system and functions of B cells, how myelin antigens emerge and present to the immune system is under-explored (*34*). Our model with vesicular impairment in oligodendrocytes proposed an endogenous pathway for disrupting myelination leading to MS, fitting in the “inside-out” model. Damages of oligodendrocytes have been observed in the early stages of MS (*35*). Genetic analyses revealed risk genes and disease-associated oligodendrocytes often with abnormal inflammation (*6-8, 36*). Nevertheless, majorities of “inside-out” models target myelin disruption or oligodendrocytes death directly without considering the mechanism of the gliopathy (*24-26, 37, 38*). Disruption of vesicular transport in oligodendrocytes caused myelin swelling and blistering, which is present in early MS lesions (*39*). Hence, oligodendrocytes are crucial in subclinical MS progression in damaging myelin.

Previous “inside-out” and oligodendrocyte-induced demyelination provided contradicting results on peripheral immune cell responses. Cuprizone-induced demyelination does not cause a T cell response unless coupled with immune stimulation (*40*). Models that induced oligodendrocytes apoptosis or cell death failed to provide immediate autoimmunity (*9, 24-26, 38*). On the contrary, disruption of transcription factors regulating lipid metabolism in oligodendrocytes without causing cell death leads to CD3+ T cells infiltration into the CNS (*12*). As such, along with our study, dysfunctional oligodendrocytes are potentially the main pathway in immune system activation. Oligodendrocytes in MS increase expression in Major Histocompatibility complex (MHC) type I and II, suggesting oligodendrocytes have the potential to present antigens to T cells (*6, 41*). Our model demonstrated abnormal accumulation of MAG and MOG in oligodendrocytes, which can potentially predispose cells to present these proteins to T cells leading to an autoimmune attack on myelin.

Remyelination was evident yet incomplete in our SNAP-23 icKO model. One of the major remyelinating strategies is the generation of new oligodendrocytes from OPCs (*21, 42, 43*). VAMP-2/3 inactivation in oligodendrocytes affects OPCs development due to impaired secretion of necessary growth factors (*20*). Oligodendrocytes are known to secrete a wide range of growth factors and cytokines (*44, 45*). The incomplete remyelination could be potentially due to altered secretion from oligodendrocytes by SNAP-23 removal. Another possibility for incomplete functional recovery was axonal damage, such as axonal spheroids. Axonal spheroids cause impaired axonal conduction and increased amyloid-beta plaque deposition (*28, 29*). Along with our studies, various animal models disrupting myelination cause amyloid precursor protein (APP) accumulation (*11, 13, 29*). Independent of myelination, oligodendrocytes express monocarboxylate transporter 1 (MCT-1) and secretes exosomes to metabolically support axonal functions and survival (*46, 47*). These functions could also be disturbed by impaired vesicular transport. Thus, insufficient support to myelinated axons from oligodendrocytes could underlie grey matter damage leading to cognitive deficits in MS.

Unexpectedly, we observed a loss of black fur color in the SNAP-23 icKO mice. Plp1-CreER^T^ can drive knockout in peripheral myelinating Schwann cells but with lower efficiency (*48*). As such, SNAP-23 can also be removed in the peripheral nervous system impacting myelination. Myelin injury can cause Schwann cells to de-differentiate into precursor cells and generate melanocytes for skin pigmentation (*32, 49*). SNAP-23 plays an essential role in transporting tyrosinase-related protein 1 (Tyrp1) in melanocytes to synthesize melanin (*50*). Hence, the white fur color might come from SNAP-23 knockout in melanocytes converted from PLP+ Schwann cells.

Impairing vesicular transport in oligodendrocytes in young adolescence could predispose demyelination with autoimmune inflammation in adults. Hence, we proposed oligodendrocyte damage as an early mechanism driving MS progression. With protein mistrafficking and altered vesicular fusion identified as molecular pathways, targeting oligodendrocytes will be a promising treatment for MS.

## Acknowledgments

The authors wish to thank the Nanoscale Biomedical Imaging Facility, The Hospital of SickKids, Toronto, Canada, for assistance with the transmission electron microscopy experiments. We acknowledge Drs. Michael Reber and Olga Rojas for providing valuable feedback on this project. We thanked the technicians and vets from the Animal Resource Centre (ARC) in the Krembil Research Institute for their help in animal care. We also thanked all the members from the Sugita Lab for their helpful discussion and inputs.

## Funding

Natural Sciences and Engineering Research Council of Canada RGPIN 2020 07139 (SS)

Canadian Institute of Health Research CIHR PJT 165917 (SS)

## Author contributions

Conceptualization: SS, CHC

Methodology: SS, CHC, MH, KO, LZ

Investigation: CHC, MH, JR, LZ, HH, SE

Visualization: CHC, MH, JR

Funding acquisition: SS, CHC, MH

Project administration: SS, CHC

Supervision: SS

Writing – original draft: CHC, SS

Writing – review & editing: SS, CHC, HS, ZF, PM, KO, LZ

## Data and materials availability

All data are available in the main text or the supplementary materials.

## Supplementary Materials

### Materials and Methods

#### Animals

Male and female mice are used in this study. The ages of the animals were specified in each experiment. C57/B6 Plp1-CreERT mice are obtained from the Jackson Laboratory (005975). SNAP-23 flox/flox mice were previously obtained (*51*). SNAP-23 flox/flox mice were crossed with Plp1-CreERT mice to generate the transgenic mouse Plp1-CreERT; SNAP-23 flox/flox. Mice were housed in numbers of one to five in a cage with food and water access ad libitum. Mice were in a room at 22-23ºC with a 12-hour light/dark cycle.

Modified EAE scoring was performed as previously described (*12*). Mice were evaluated weekly for demyelination and neurological deficits. An 8-point scale was used to investigate 4 categories, with each category having a score of 0-2. Tremor: no tremor (0), tremor activity when the mouse was held with the tail (1), tremor when walking and standing (2). Limp tail: The tail was upright and spinning when holding the base of the tail (0), the tip of the tail was flaccid (1), completely flaccid tail (2). Hind limb weakness: normal (0), the angle between the hind limb and paw was around 90º (1), the angle between the hind limb and paw was around 180º (2). Hind limb paresis: normal movement and gait, can grip wire bars firmly (0), slips through bars when walking on a grid surface, abnormal gait (1), complete hind limb paralysis (2).

#### Mouse Genotyping

A mouse tail sample was collected from a 3-week-old mouse for genotyping polymerase chain reaction (PCR). Tail samples were dissolved in 50mM NaOH at 95ºC for 1 hour. PCR was used to detect the presence of CreERT and SNAP-23 flox allele. For Plp1-CreERT, the forward primer is 5’-ATACCGGAGATCATGCAAGC-3’ and the reverse primer is 5’-GGCCAGGCTGTTCTTCTTAG-3’. For the SNAP-23 flox allele, the forward primer is 5’-GGGGGTGAGTTGAAGTCATTGAAG-3’ located at the end of the exon 4, the reverse primer is 5’-AGCTTAAACGGGATGAACTCAGGC-3’ located at the start of the intron 4.

#### Tamoxifen preparation and injection

Tamoxifen was administered to 3-to-4-week-old mice at 100mg/kg body weight. Tamoxifen was first dissolved in ethanol at 20mg/mL, then layered on an equal amount of corn oil (Sigma Aldrich, CAS: 8001-30-7). Tamoxifen was dissolved in the corn oil by spinning in a speed vacuum for 1 hour. Tamoxifen was injected into mice via intraperitoneal injection.

#### Rotarod

Mice were subjected to the accelerating rotarod for four consecutive days using the Panlab apparatus. The speed of the rotarod started from 4 rotations per minute (rpm) and reached 40 rpm in 5 minutes. The time and speed of mice falling off the rotarod were recorded.

#### Mouse tissue collection for histology and genotyping

Mice were anesthetized by intraperitoneal injection of sodium pentobarbital (75mg/kg body weight, Bimeda-MTC). Mice were then transcardially perfused with 10mL PBS followed by 10mL 4% paraformaldehyde (PFA). The brain, spinal cord, and optic nerves were dissected. The brain was fixed in 4% PFA overnight, while the spinal cord and optic nerves were fixed in 4% PFA for 2 hours. Tissues were dehydrated in 30% sucrose dissolved in PBS overnight. Brain and spinal cord samples were mounted with a solution of 50% OCT and 15% sucrose and cryosectioned at 40μm. Optic nerves were mounted in OCT solution and cryosectioned at 14μm.

#### Genotyping for knockout allele

To genotype for the SNAP-23 knockout allele, 10-15 fixed 40μm spinal cord sections were dissolved in 50mM NaOH at 95ºC for 1 hour. The forward primer, 5’-TCCAACCCAGAGAGACACTTTT-3’, located upstream of exon 3 and the reverse primer, 5’-GGTACTGTTGCCTCTCATCCCAGT-3’, located downstream of exon 4 were used.

#### Luxol Fast Blue staining

Cryosectioned tissues were placed in 0.1% LFB solution (Thermo Scientific, CAS: 1328-51-4) at 56ºC overnight. Sections were differentiated with 0.05% Lithium carbonate solution (TCI America, CAS: 554-13-2) and 70% alcohol until grey and white matters were differentiated. After washing with distilled water, sections were mounted with cytoseal 60 (Epredia). Quantification of LFB staining was done in Fiji (ImageJ). LFB positive area was determined by auto thresholding and compared with the original image.

#### Immunohistochemistry

Sectioned samples were washed and permeabilized with 0.1% Triton X-100 in PBS and blocked with 10% goat serum. Samples were incubated in the primary antibody at room temperature overnight. Primary antibodies used: rat anti-MBP (1:50, Millipore Sigma, MAB386), rabbit anti-PLP (1:500, Abcam, ab284363), rabbit anti-MAG (1:100, Cell Signaling Technology, 9043), rabbit anti-MOG (1:500, Abcam, ab32760), mouse anti-OLIG2 (1:500, Millipore Sigma, MABN50), rabbit anti-GFAP (1:100, Cell Signaling Technology, 12389), rabbit anti-IBA1 (1:100, Cell Signaling Technology, 17198), rabbit anti-CASPR (1:500, Abcam, ab34151), mouse anti-Ankyrin G (1:250, Millipore Sigma, MABN466), and rat anti-CD3 (1:100, Invitrogen, 14-0032-82). The next day, sections were washed with 0.1% Triton X-100 in PBS again before staining with DAPI and the secondary antibody that matched the species of the primary antibody for 1 hour. Samples were then subjected to another 0.1% Triton X-100 in PBS wash. If samples were to stain with FluoroMyelin, slides with samples were flooded with 1:300 diluted FluoroMyelin Red (Invitrogen) for 20 minutes. Sections were mounted in Mowiol. For confocal imaging, Carl Zeiss confocal microscope was used. Images were taken at oil immersion 40X objective at 1.0 Airy Unit pinhole sizes. Zeiss immersion oil 518F was used. Two-photon imaging was performed with a two-photon microscope (Custom made FV1000 MPE; Olympus, Tokyo, Japan, 60x objective lens, NA 1.0 equipped with Spectra-Physics InSight DeepSee; Spectra-Physics, CA, US). The fluorescence images were prepared from the maximum intensity projection across a z-stack, taken at 1 μm intervals by Fluoview (Olympus, Tokyo, Japan).

#### Transmission Electron Microscopy

Optic nerves transmission electron microscopy (TEM) was performed as previously described (*52*). Mice were anesthetized as above. After transcardial perfusion of 10mL PBS, mice were perfused with 10mL 2% glutaraldehyde (in 0.1M sodium cacodylate buffer, pH 7.3). Optic nerves were dissected and cut into 2mm sections and fixed with 2% glutaraldehyde overnight. Tissues were post-fixed with 1% osmium tetroxide and dehydrated with alcohol. Tissues were infiltrated with Spurr resin and polymerized at 65ºC overnight. Tissue blocks were then sectioned at 90nm with an ultramicrotome and collected on 200 mesh copper grids. Sections were stained with uranyl acetate and lead citrate before imaging with the Hitachi HT7800 TEM.

#### *Ex vivo* compound action potential recording of optic nerves

Mice were anesthetized as above. Mice were transcardially perfused with cold dissection solution (300 mM sucrose, 3.5 mM KCl, 2 mM NaH_2_PO_4_, 20 mM glucose, 0.5 mM CaCl_2_, 2 mM MgCl_2_ and 5 mM HEPES (pH adjusted to 7.4)). The optic nerves were dissected and placed in artificial cerebrospinal fluid (ACSF, 125 mM NaCl, 25 mM NaHCO_3_, 3.5 mM KCl, 1 mM NaH_2_PO_4_, 1 mM MgSO_4_, 2 mM CaCl_2_) with oxygenation (95% O_2_ and 5% CO_2_) at room temperature at least 3 hours before recording.

Whole optic nerve compound action potential was recorded as previously described (*53*). For recording, optic nerves were placed in 37ºC oxygenated ACSF. Two suction electrodes (A-M System, Sequim, WA USA were placed on the two ends of the optic nerves. A stimulating electrode was placed on the retinal end and a recording electrode was at the chiasmal end. Syringes attached to the electrodes were used to create suction at both ends without stretching the nerves. The stimulus was delivered in a 1ms pulse with an amplitude that produced the maximal intensity of CAP. CAP traces were an average of 20 trials. Tetrodotoxin (1 μM) was applied to block CAP conduction to obtain traces of stimulus artefacts. CAP traces minus stimulus artefacts obtained pure CAP waveform for analysis. Signals were recorded with a dual channel amplifier (700B, Molecular Devices/ Axon Instruments, Sunnyvale, California, USA) and digitized at 10kHz (Digidata 1322A, Molecular Devices). Data was acquired with PClamp software (version 10, Molecular Devices). Stimulus artefact removed CAP data was analyzed in OriginLab (2022). CAP traces were fitted with three Gaussian functions to determine peak amplitude, latency, and full width at half-maximum.

#### Visual evoked potential recording

The Visual evoked potential (VEP) was recorded as previously described (*21*). Mice were anesthetized with Avertin (200mg/kg) and kept in the dark for 5 minutes. Mydriacyl (Alcon) eye drops were applied to dilate pupils in both eyes. A steel recording electrode was placed subcutaneously at the visual cortex at 8mm in depth. A ground electrode was placed at the tail and a reference electrode was placed between the eyes subcutaneously. White light stimulation (6500K) at pulse intensity 3cd.s/m2, frequency at 1Hz, and duration at 4ms was delivered using the Diagnosys Espion electroretinogram device with a ColorDome Ganzfeld stimulator and Espion software. A total of three runs were performed per mouse with each run containing 100 sweeps of light stimulation. Body temperature was kept at 38ºC throughout. VEP for analyzing remyelination in mice was performed similarly except mice were anesthetized with Pentobarbital (60mg/kg body weight).

#### Statistical analysis

Statistical analysis was performed in RStudio (Posit, PBC) and OriginLab 2022 (OriginLab, Northampton, MA). All data are mean±SEM. Statistical tests performed were specified in each experiment. p<0.05 was considered statistically significant in all experiments. For box plots, the central line defined the median, box boundaries defined the 25th and 75th percentile, and whiskers defined the full range of data.

## Figs S1 to S7

**Fig. S1.**
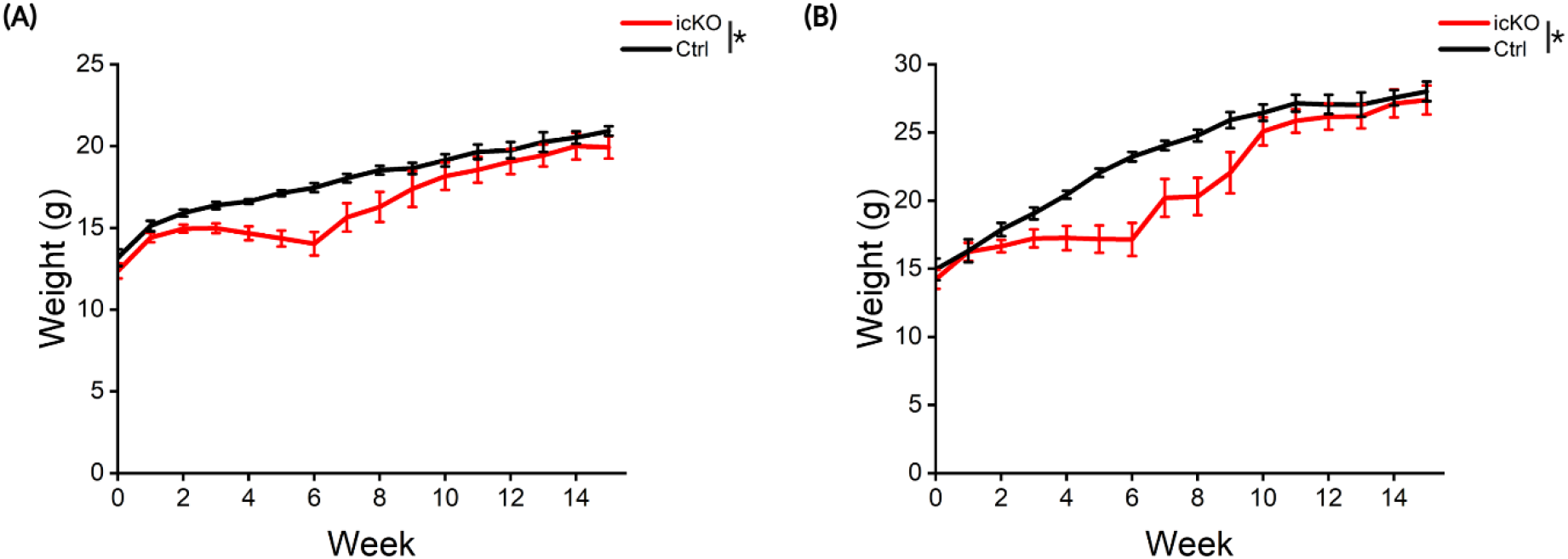
Weight decline with demyelination in SNAP-23 icKO. Weight loss in females (ctrl: n=4-13; SNAP-23 icKO: n=6-18) **(A)** and males (ctrl: n=6-10; SNAP-23 icKO: n=6-13) **(B)** after tamoxifen injection. Data are mean±SEM. *p<0.05 by 2-way ANOVA followed by Tukey post hoc test.

**Fig. S2.**
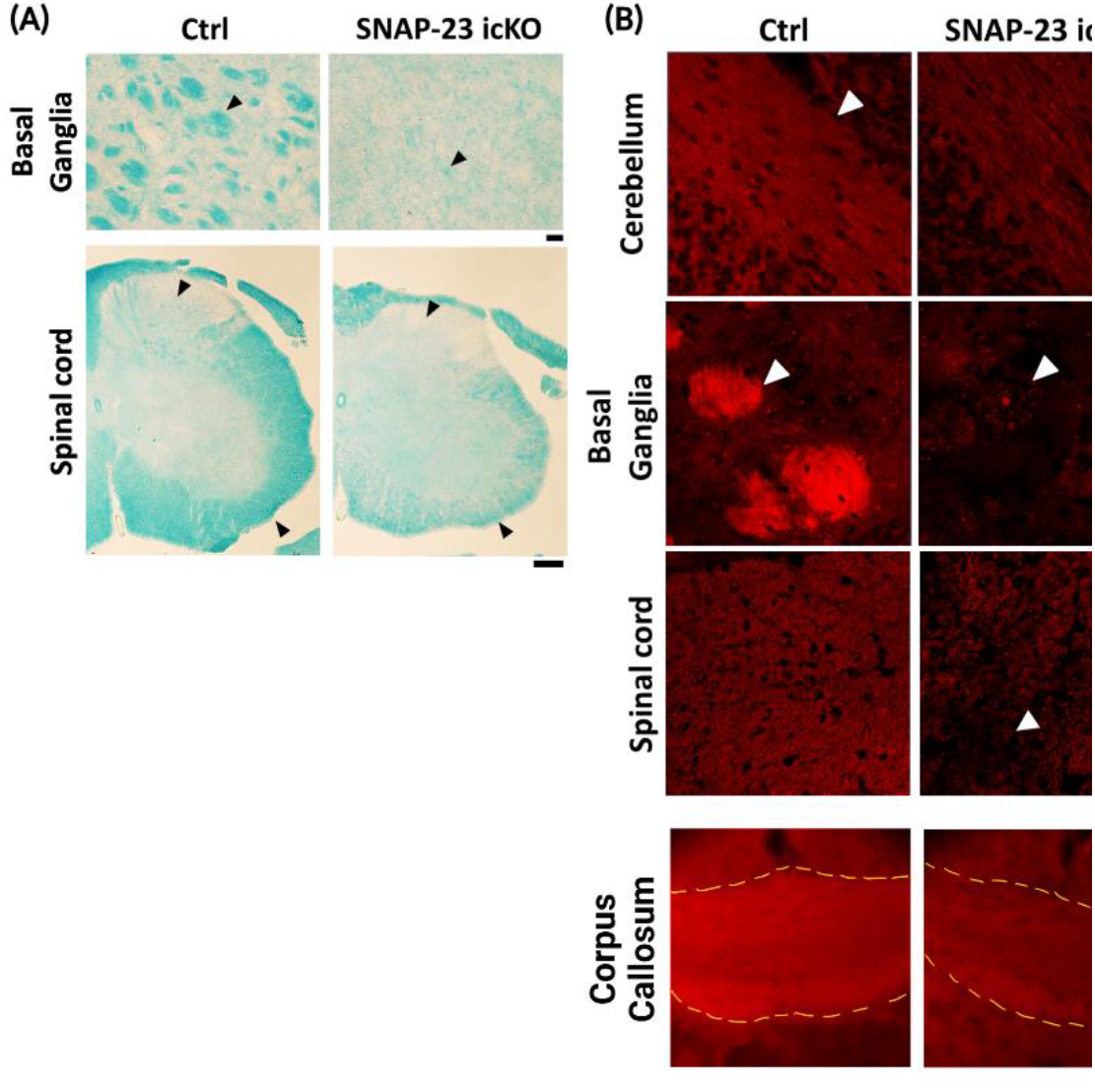
Demyelination in the CNS by LFB and FluoroMyelin staining. **(A)** LFB staining of the basal ganglia (Scale bar: 50μm) and the spinal cord (Scale bar: 500μm). Arrowheads indicate myelin regions to compare. **(B)** Demyelination was verified by FluoroMyelin staining in the cerebellum, basal ganglia, and spinal cord (indicated by white arrowheads, scale bar: 20μm). Fluoromyelin staining was also reduced in the corpus callosum (Scale bar: 40μm).

**Fig. S3.**
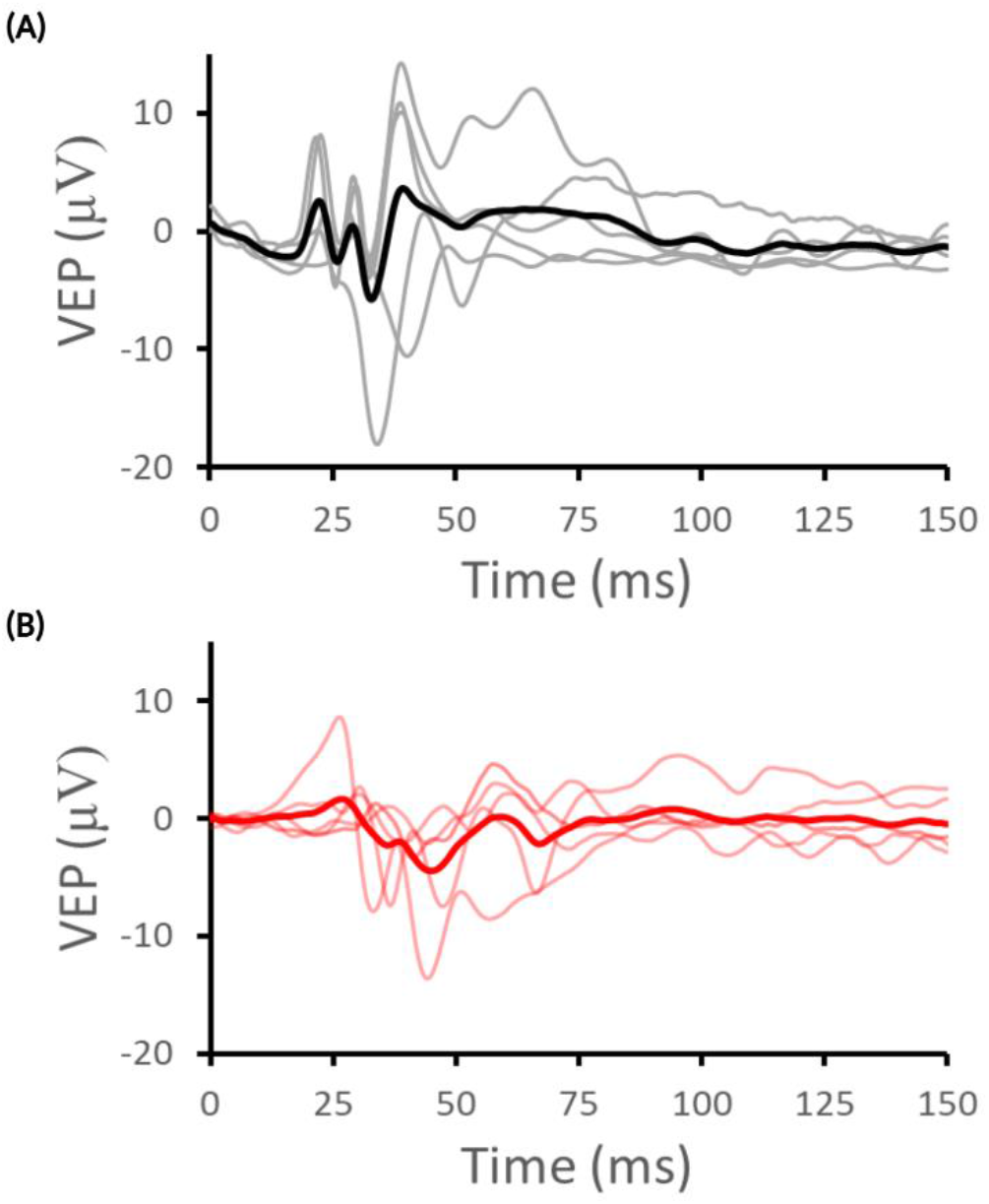
Individual VEP traces. VEP traces of ctrl (n=5) **(A)** and SNAP-23 icKO (n=5) **(B)**. Dark color traces represent the average traces of each group.

**Fig. S4.**
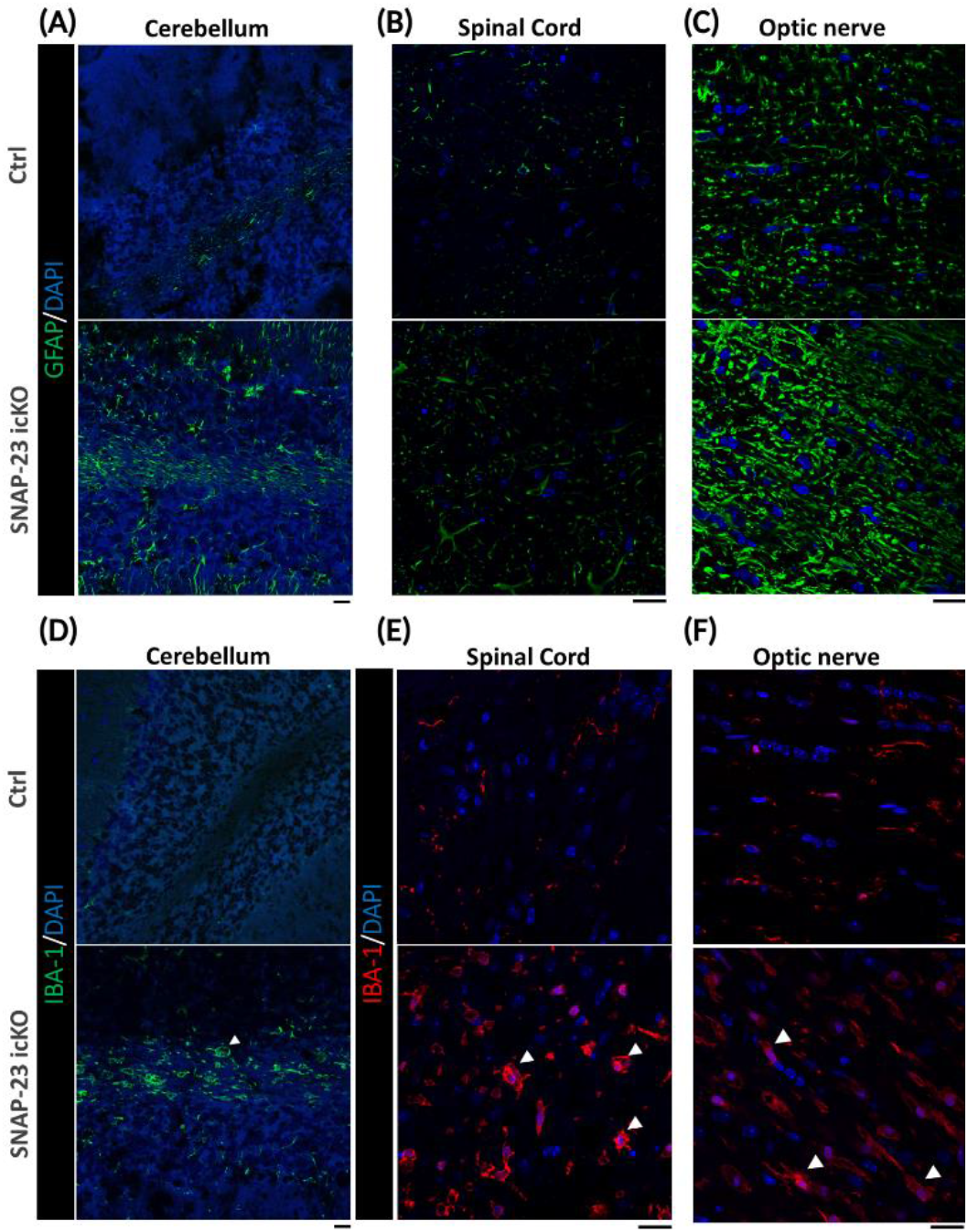
Neuroinflammation across different regions in the CNS. **(A-C)** GFAP staining of astrocytes in the cerebellum (A), spinal cord (B), and optic nerve (C). **(D-F)** IBA-1 staining of microglia in the same regions as above. Arrowheads indicate microglia. Scale bar: 20μm.

**Fig. S5.**
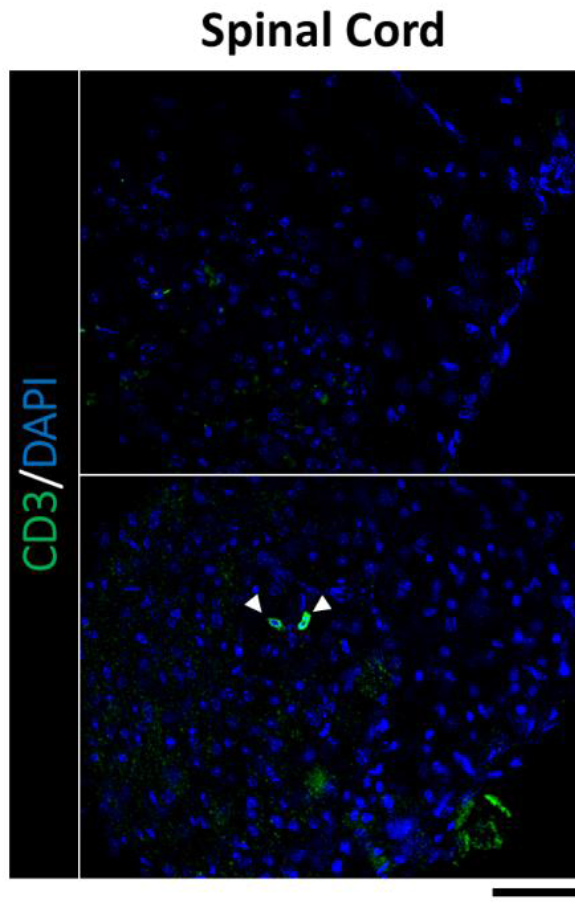
CD3 T cells infiltration into the spinal cord. Arrowheads indicate CD3+ cells. Scale bar: 20μm.

**Fig. S6.**
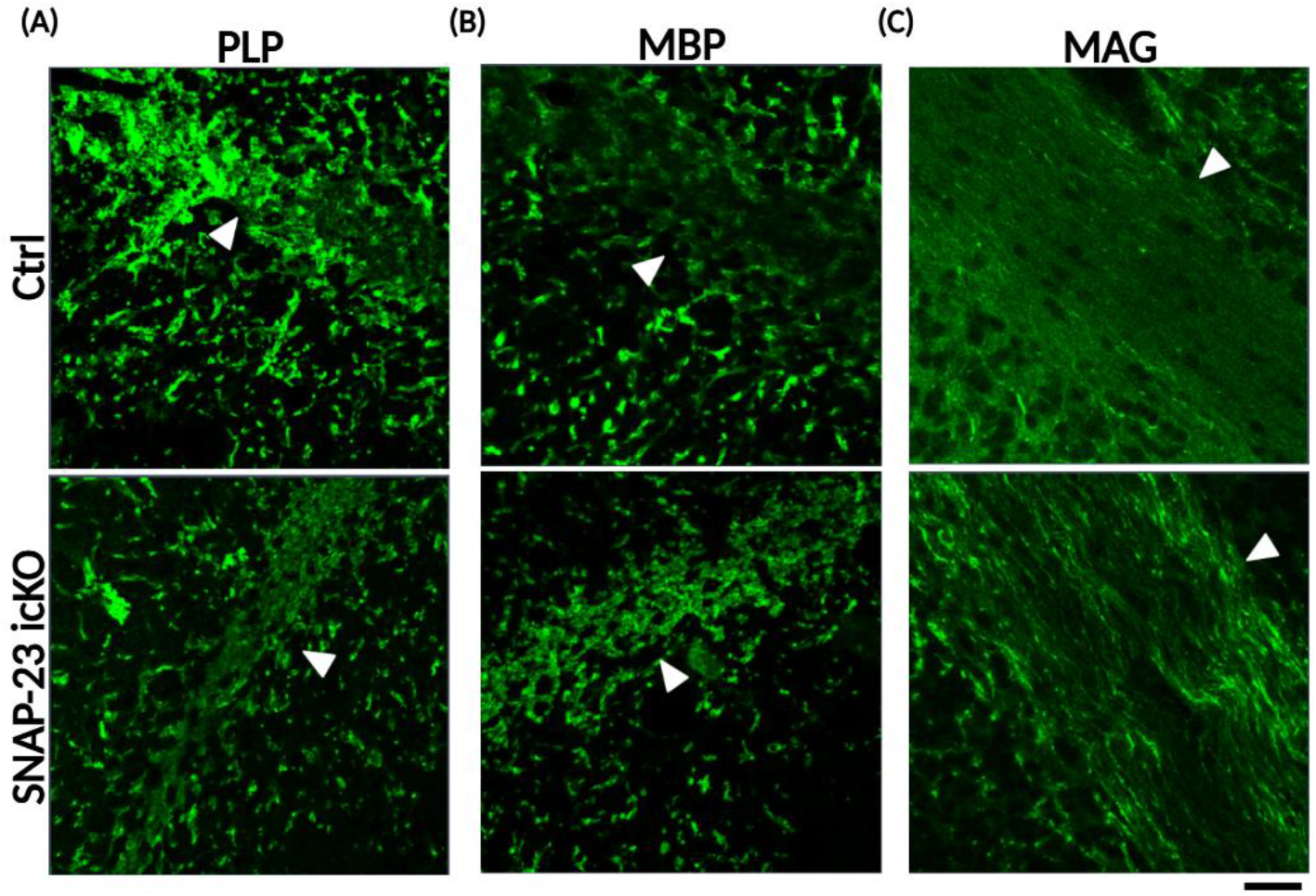
SNAP-23 removal impaired MAG trafficking with little effects on PLP and MBP in oligodendrocytes. **(A-C)** Immunostaining of PLP (A), MBP (B), and MAG (C) in the cerebellum (White arrowhead pointing to the myelin tract). Scale bar: 20μm.

**Fig. S7.**
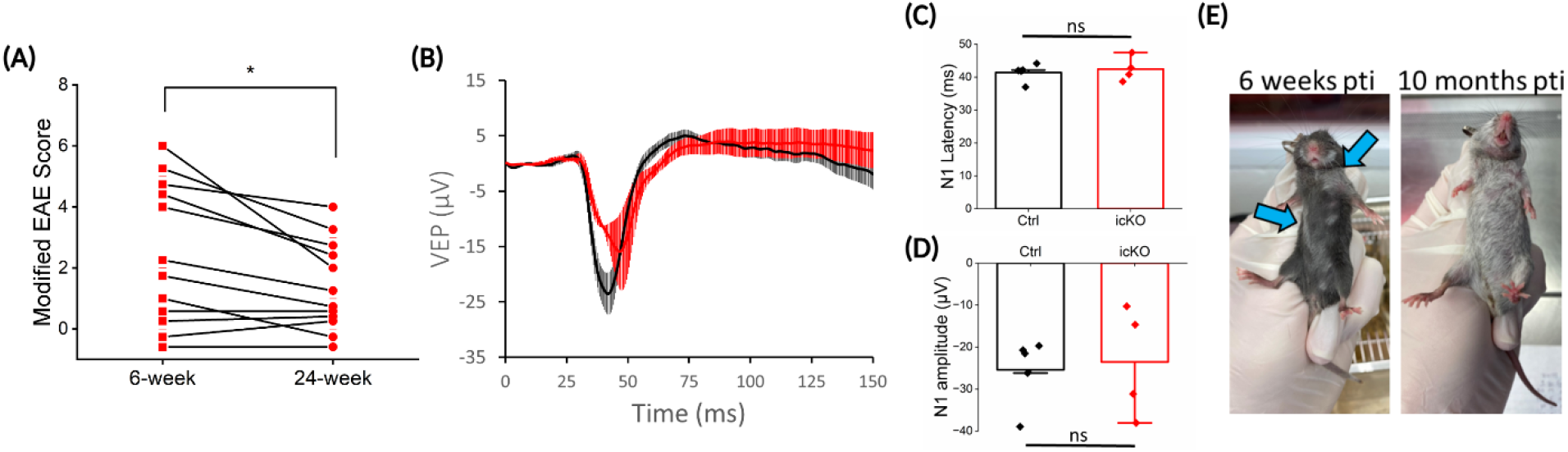
Remyelination in SNAP-23 icKO after demyelination. **(A)** Reduction in modified EAE score observed 24 weeks (6 months) pti compared to 6 weeks (peak demyelination). *p<0.05 by pair sample t-test. **(B-D)** VEP measurement in ctrl (n=5) and SNAP-23 icKO (n=4) 2 to 7 months pti. No difference in N1 latency and amplitude (two-sample t-test). Data are mean±SEM. **(E)** White fur color in 6 weeks pti (indicated by arrows) and 10 months pti. The whole mouse changed into white color in 10 months pti.

